# Chitin biosynthesis genes in *Diaphorina citri*, Asian citrus psyllid

**DOI:** 10.1101/2020.09.22.309211

**Authors:** Sherry Miller, Teresa D. Shippy, Blessy Tamayo, Prashant S Hosmani, Mirella Flores-Gonzalez, Lukas A Mueller, Wayne B Hunter, Susan J Brown, Tom D’elia, Surya Saha

## Abstract

The polysaccharide chitin is critical for the formation of many insect structures, including the exoskeleton, and is required for normal development. Here we report the annotation of three genes from the chitin synthesis pathway in the Asian citrus psyllid, *Diaphorina citri* (Hemiptera: Liviidae), the vector of Huanglongbing (citrus greening disease). Most insects have two chitin synthase (CHS) genes but, like other hemipterans, *D. citri* has only one. In contrast, *D. citri* is unusual among insects in having two UDP-N-acetylglucosamine pyrophosphorylase (UAP) genes. One of the *D. citri* UAP genes is broadly expressed, while the other is expressed predominantly in males. Our work helps pave the way for potential utilization of these genes as pest control targets to reduce the spread of Huanglongbing.

## Introduction

Chitin is a polysaccharide essential for insect development. It plays a crucial role in the development of the insect cuticle and exoskeleton, the peritrophic membrane of the midgut of some insects, and other structures such as the trachea, wing hinges and eggshell [1]. The biosynthetic pathway for chitin involves several enzymes that act on simple sugars such as glucose, trehalose and glycogen to produce intermediates that are subsequently converted into chitin. In the penultimate step of the chitin biosynthesis pathway, N-acetylglucosamine-1- phosphate is converted into UDP-N-acetylglucosamine. This reaction is catalyzed by the enzyme UDP-N-acetylglucosamine pyrophosphorylase (UAP) [1]. In the final step of the pathway, UDP-N-acetylglucosamine is converted to chitin by enzymes known as chitin synthases (CHS) [1]. Because chitin is essential for insect development, but is not found in mammals, the enzymes involved in its synthesis are considered attractive targets for pest control. Here we report the annotation of one CHS gene and two UAP genes in the Asian citrus psyllid, *Diaphorina citri* (Hemiptera: Liviidae), the vector for the bacterium that causes Huanglongbing (citrus greening disease). Although most insects have two CHS genes [2,3] (Table 1), the presence of a single CHS gene is consistent with reports from other hemipteran genomes [4]. In contrast, *D. citri* seems to be unusual in having two UAP genes. RNA-Seq data indicates that one of the *D. citri* UAP genes is broadly expressed, while the other is expressed predominantly in males. Our manual annotation of these chitin biosynthesis genes provides improved gene targets for future experiments.

**Table 1.** Gene counts are taken from published reports [2,4,20] or determined from genome data [29–36]. *D. citri* numbers are based on annotation of genome v3.

|  | *Drosophila*<br>*melanogaster* | *Anopheles*<br>*gambiae* | *Aedes*<br>*aegypti* | *Tribolium*<br>*castaneum* | *Apis*<br>*mellifera* | *Nasonia*<br>*vitripennis* | *Acyrtosiphon*<br>*pisum* | *Bemisia*<br>*tabaci* | *Diaphorina*<br>*citri* |
| --- | --- | --- | --- | --- | --- | --- | --- | --- | --- |
| CHS1/A | 1 | 1 | 1 | 1 | 1 | 1 | 1 | 1 | 1 |
| CHS2/B | 1 | 1 | 1 | 1 | 1 | 1 | 0 | 0 | 0 |
| UAP | 1 | 1 | 1 | 2 | 1 | 1 | 1 | 1 | 2 |

## Results and Discussion

### Chitin Synthases

Chitin synthases are the only enzymes in the chitin biosynthetic pathway that act specifically in the synthesis of chitin, making them an attractive, insect-specific target for RNA interference (RNAi) based insecticides. The two *CHS* genes found in most holometabolous insects have distinct functions. *CHS1*, also referred to as *CHSA*, produces the chitin essential for proper cuticle development [3,5,6]. *CHS2*, also referred to as *CHSB*, is not required for cuticle development, but is instead essential for proper development of the gut peritrophic membrane [3,5,6]. RNAi knockdown of either *CHS gene* is lethal in holometabolous insects [7–10].

Previous searches of the *Acyrthosiphon pisum, Nilaparvata lugens* and *Rhodnius prolixus* genomes identified *CHS1* but not *CHS2*, suggesting that *CHS2* has probably been lost in the hemipteran lineage [4]. Loss of the chitin synthase gene required for peritrophic membrane development is not particularly surprising, since hemipterans do not have peritrophic membranes [4,11]. Lu et al [12] identified a *D. citri CHS* gene that clustered with other hemipteran CHS genes and was expressed at high levels in most adult body tissues, but at low levels in midgut, as would be expected for a *CHS1* gene. Two groups have shown that RNAi knockdown of *CHS* in *D. citri* causes increased lethality [12,13], supporting the idea that this gene is a good target for pest control.

Our searches of the *D. citri* v3 genome revealed the previously described *CHS* gene, but no additional chitin synthase orthologs (Table 1). Transcriptomic evidence supports the existence of two *CHS* isoforms (Table 2) that differ only in the use of one alternative exon and produce proteins with slightly different C-termini. Similar isoforms of *CHS1/A* have been described in other insects [2,14,15]. Both isoforms of *D. citri* CHS cluster in a monophyletic clade with CHS1 proteins from other insects (Figure 1), so we have named this gene *CHS1. We* retrieved expression data for both isoforms of CHS1 from the Citrusgreening Expression Network (CGEN), which contains RNA-Seq data sets for a variety of life stages and tissues [16]. Data from whole body samples indicate that CHS1 is expressed at all life stages, but is most highly expressed in juvenile stages (Figure 2).

**Table 2.** Each manually annotated gene is full length and has been assigned an OGSv3 gene identifier. Evidence types used for manual annotation are shown for each gene. A description of the various evidence sources and their strengths and weaknesses is included in our online protocol [25].

| Gene/Isoform | OGSv3 ID | Evidence supporting annotation |  |  |  |
| --- | --- | --- | --- | --- | --- |
|  |  | MCOT | IsoSeq | RNASeq | Ortholog |
| CHS1 | Dcitr04g09970.1.1<br>Dcitr04g09970.1.2 | X | X | X | X |
| UAP1 | Dcitr08g04630.1.1 |  | X |  | X |
| UAP2 | Dcitr05g05060.1.1 |  | X | X | X |

**Figure 1:**
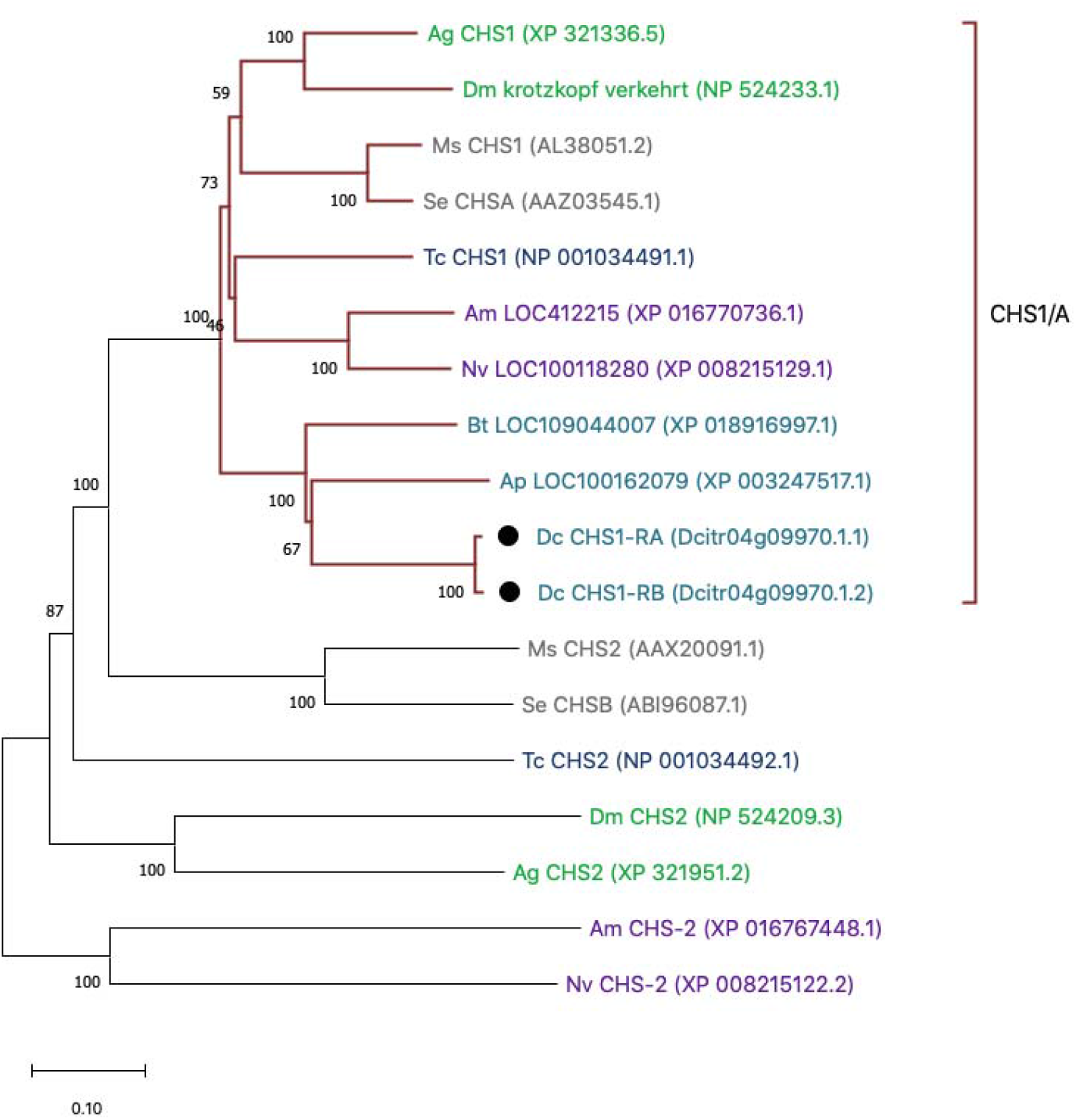
Phylogenetic analysis of insect CHS proteins. Species represented are *Drosophila melanogaster* (Dm), *Anopheles gambiae* (Ag), *Tribolium castnaeum* (Tc), *Manduca sexta* (Ms), *Spodoptera exigua* (Se), *Apis mellifera* (Am), *Nasonia vitripennis* (Nv), *Acyrthosiphon pisum* (Ap), *Bemisia tabaci* (Bt) and *Diaphorina citri* (Dc). MUSCLE software was used to perform multiple sequence alignments of full-length protein sequences and the tree was constructed with MEGA X software using the neighbor-joining method with 100 bootstrap replications. The maroon clade shows monophyletic clustering of CHS1/A genes. With the exception of *D. citri* (denoted by black circles), only one isoform per species is depicted. Taxon name color represents insect order: Diptera (green), Coleoptera (navy), Hymenoptera (purple), Lepidoptera (gray), and Hemiptera (teal).

**Figure 2.**
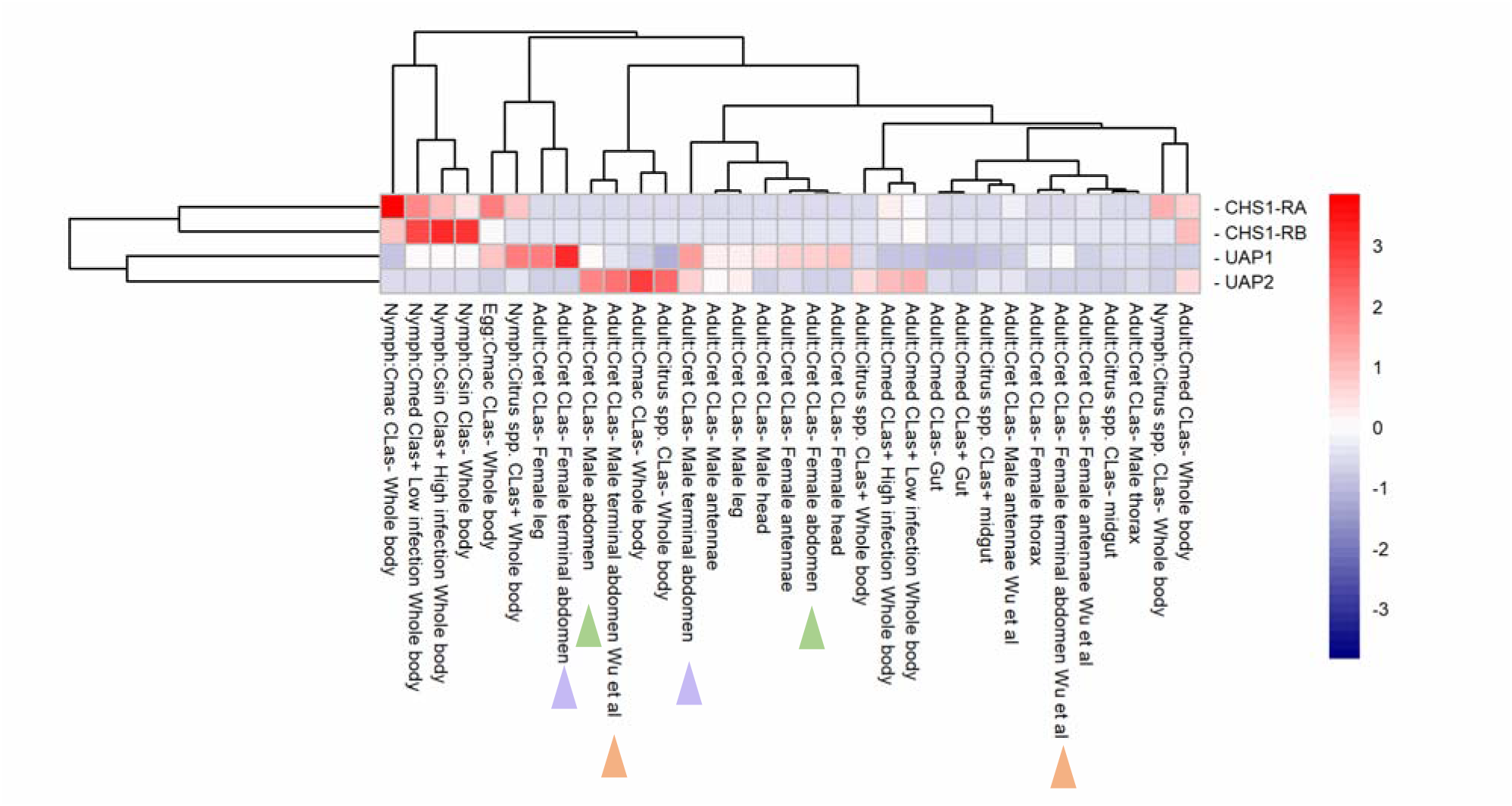
Heatmap representation of chitin biosynthesis gene expression levels in a variety of RNA-Seq datasets. Expression levels were obtained as transcripts per million (TPM) from the Citrusgreening Expression Network [16]. For ease of comparison, colored arrowheads denote pairs of male and female abdominal tissue samples from the same experiments.

Our manual annotation of *CHS1* corrects several errors that were present in the previous computationally-predicted annotation for *D. citri* CHS (XP_017303059). Changes to the model include the addition of formerly missing sequence and the removal of artifactually duplicated regions. Domain analysis with TMHMM Server, v. 2.0 indicates that the corrected CHS1-RA and CHS1-RB proteins have 15 transmembrane helices as is typical for insect CHS proteins, rather than the 14 that were reported for the earlier version of the protein [12].

### UDP-N-acetylglucosamine pyrophosphorylase (UAP)

In addition to its role in chitin synthesis, UAP is also involved in the modification of other carbohydrates, sphingolipids and proteins. In Drosophila, mutants of *UAP* (also called *mummy, cabrio* and *cystic)* have defects in tracheal development, dorsal closure, eye development and nervous system function [17–19]. Some of these developmental defects are due to disruption of the chitin synthesis pathway, while others appear to be caused by effects on other glycoproteins. For example, defects in embryonic dorsal closure have been linked to a role for UAP in regulation of Decapentaplegic signaling [20].

Most insects appear to have a single *UAP gene* [21]. However, a few insects, including *T. castaneum, Locusta migratoria* and *Leptinotarsa decemlineata* have two UAP genes [21–23]. Comparison of the *T. castaneum* and *L. migratoria* gene pairs indicates that they arose through separate, relatively recent lineage-specific gene duplications [22]. RNAi experiments in *T. castaneum* showed that UAP1 is involved in the biosynthesis of chitin both in the cuticle and the peritrophic membrane, while UAP2 is important for the modification of other macromolecules [21]. In *L. migratoria, LmUAP1* knockdown caused lethality and defects consistent with disruption of chitin biosynthesis, while *LmUAP2* knockdown did not increase lethality and produced no visible effects [22].

In the *D. citri* v3 genome, we identified two *UAP* genes located on different chromosome-length scaffolds. The proteins encoded by these apparent paralogs share 50 percent identity distributed throughout the length of the proteins (Figure 3), which is very similar to the level of identity shared with UAP orthologs from closely related insect species. Amino acid residues known to be important for substrate binding in the human UAP ortholog and conserved in the *T. castaneum* UAP proteins [20] are also well conserved in the *D. citri* UAP proteins (Figure 3). Phylogenetic analysis (Figure 4) suggests that the two genes represent a lineage specific duplication. Surprisingly, the *D. citri* UAP proteins do not cluster with the other hemipteran UAP proteins and instead appear as an outgroup to all the other insect UAP proteins, likely suggesting the *D. citri UAP genes* are diverging rather rapidly. We have named the *D. citri* genes *UAP1* and *UAP2*, but this should not be taken to imply direct orthology with duplicated UAP genes in other insects.

**Figure 3.**
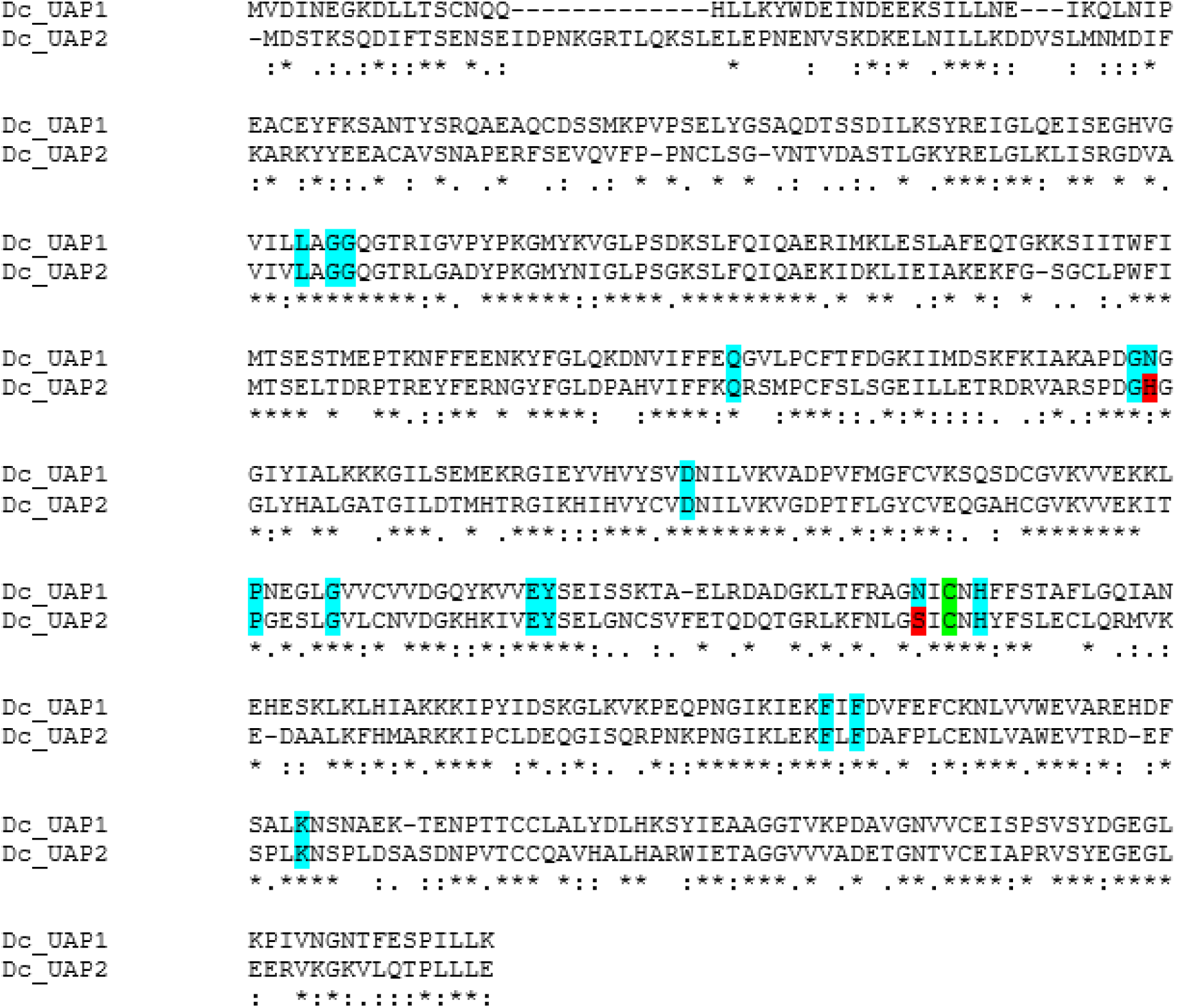
Alignment of D. citri UAP1 and UAP2. Alignment was performed using MUSCLE [25]. Individual amino acid alignments are denoted as identical (*), highly similar (:) or similar (.). Residues important for substrate binding by human UAP1 and conserved in *T. castaneum* are shaded according to their level of conservation. Identical residues are shaded blue and non-identical (but similar) residues are shaded red. The green shaded residue denotes the position of an alanine important for substrate binding in human UAP1 that is a cysteine in *T. castaneum* and other insects.

**Figure 4.**
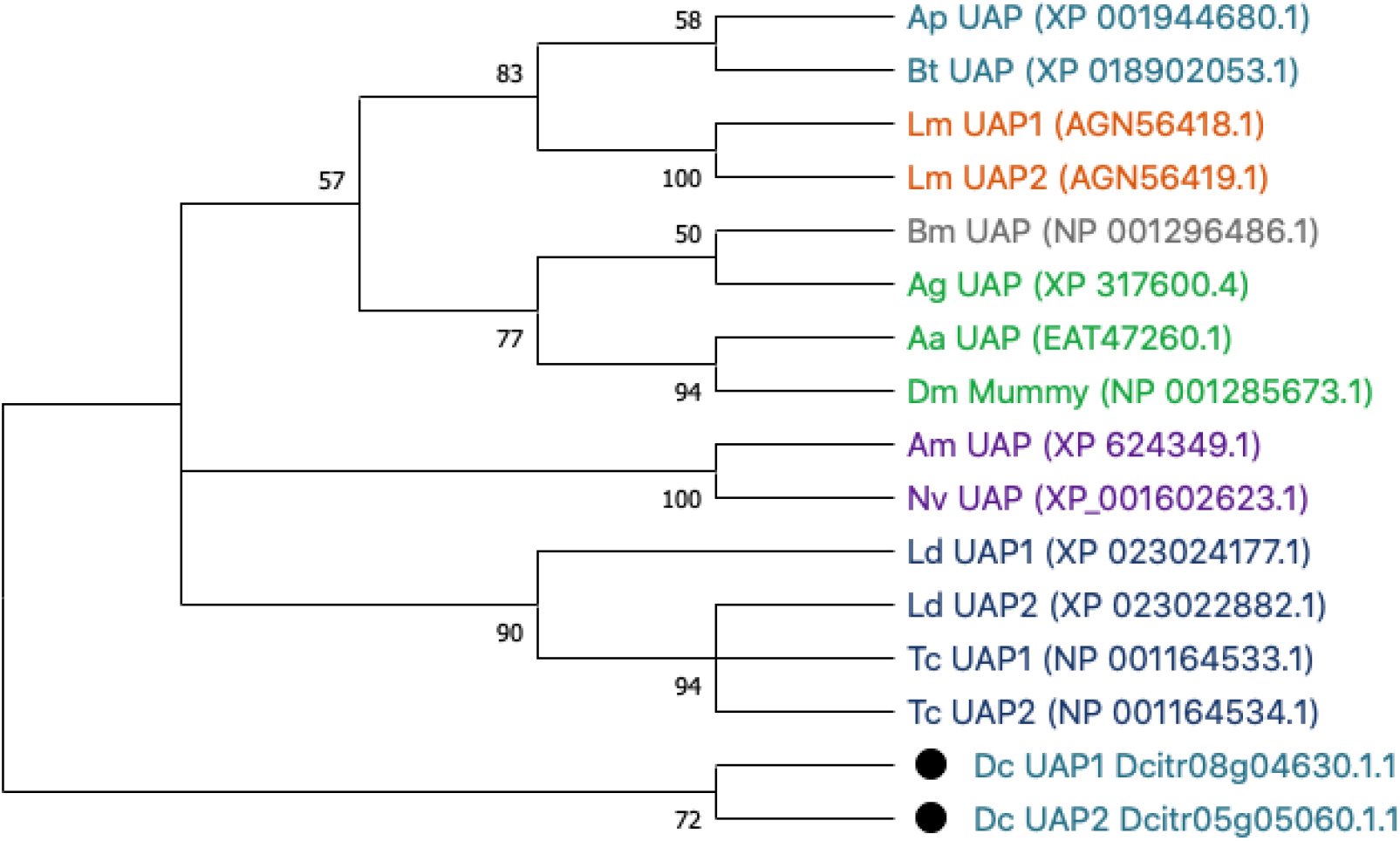
Phylogenetic analysis of representative insect UAP orthologs. Species shown are *Drosophila melanogaster* (Dm), *Anopheles gambiae* (Ag), *Aedes aegypti* (Aa), *Bombxy mori* (Bm), *Tribolium castaneum* (Tc), *Leptinotarsa decemlineata* (Ld), *Apis mellifera* (Am), *Nasonia vitripennis* (Nv), *Locusta migratoria* (Lm), *Acyrthosiphon pisum* (Ap), *Bemisia tabaci* (Bt) and *Diaphorina citri* (Dc and black circles). ClustalW software was used to perform the multiple sequence alignment of full-length protein sequences and a bootstrap consensus tree was constructed with MEGA X software using the neighbor-joining method with 100 bootstrap replications. Colors denote insect orders: Hemiptera (teal), Orthoptera (orange), Lepidoptera (gray), Diptera (green), Hymenoptera (purple) and Coleoptera (navy).

We compared available expression data from the two *D. citri UAP* genes using CGEN [16]. *D. citri UAP1* is expressed in all tissues and stages examined, although expression levels vary (Figure 2). A few samples (e.g. female terminal abdomen and female leg) show high expression of *UAP1*, but these are single replicate samples that would need further verification. In the case of female terminal abdomen, single replicate data from a separate experiment shows only a moderate level of expression. Interestingly, *D. citri UAP2* appears to show a sexually dimorphic expression pattern. It is expressed at a low to moderate level in most male tissues,with highest expression in abdominal samples, but shows little or no expression in the same tissues from females (Figures 2,5). While these observations are intriguing, the technical difficulty of creating RNA-Seq libraries from miniscule amounts of dissected tissue while maintaining integrity of the RNA in addition to the lack of statistical power provided by single replicate samples mean that the expression data currently available should be interpreted with caution. More detailed analysis of *UAP1* and *UAP2* expression and function in individual males and females will be necessary to resolve these questions.

**Figure 5.**
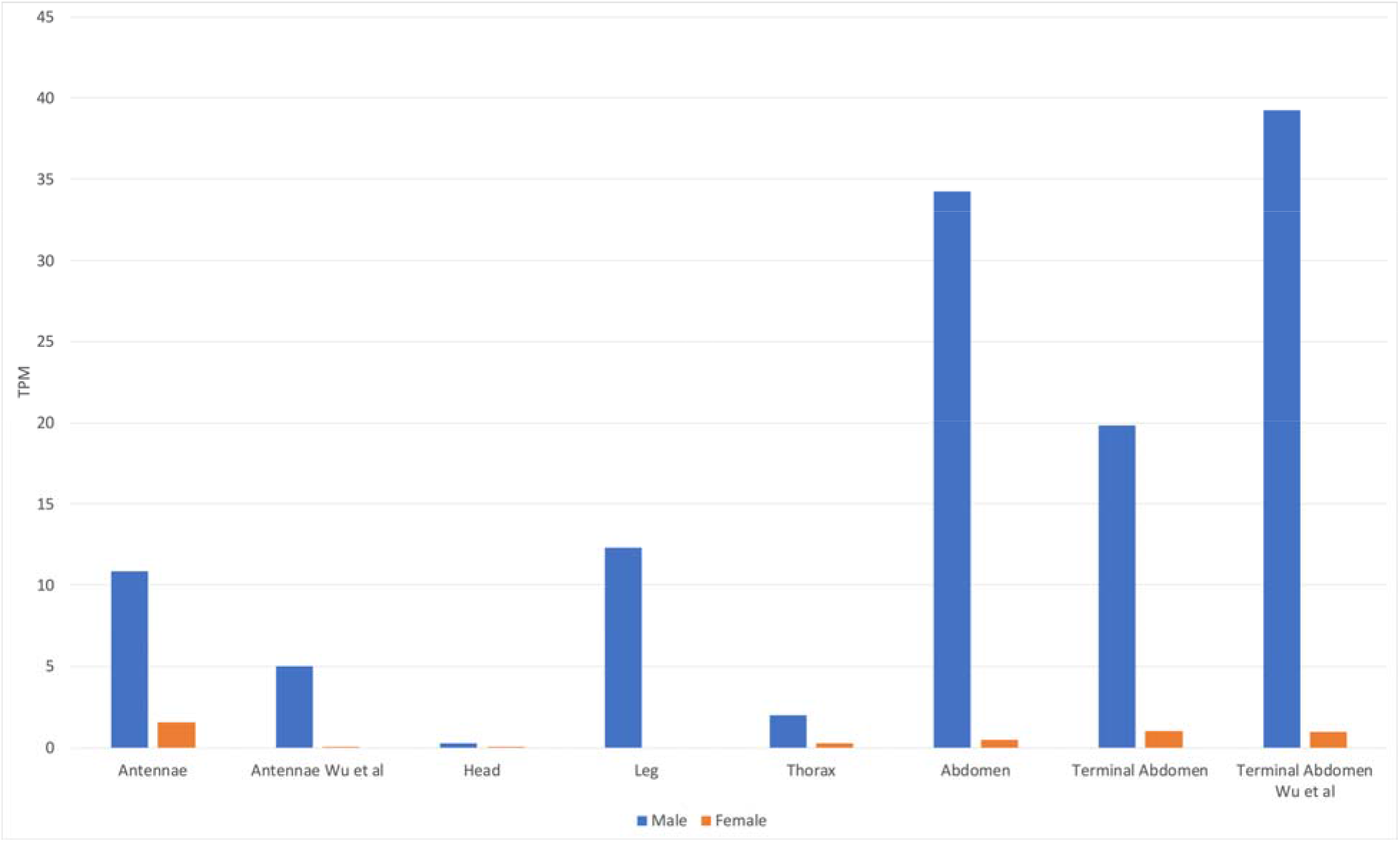
Expression levels of *UAP2* in male and female tissues. Expression levels were obtained from the Citrusgreening Expression Network [16]. Tissue types are shown on the X axis and expression levels (TPM) on they Y-axis. Blue bars denote expression levels in males and orange bars denote expression levels in females (all single replicate data). RNA-Seq data from tissues labeled Wu et al were sequenced in [37]. Data for the remaining tissues are from NCBI BioProject PRJNA448935.

### Conclusion

We searched for orthologs of 33 Drosophila segmentation genes in the *D. citri* v3 genome and identified and annotated 24 homologous genes. We were unable to find orthologs for 10 of the *Drosophila* genes, while *D. citri* has one segmentation gene (*otd-2*) whose ortholog has been lost in *Drosophila.* Most of these absences, except *eagle* and *invected*, were expected based on the known phylogenetic distribution of the genes. While all the genes discussed in this report were initially identified because of their role in embryonic patterning and segmentation in *Drosophila*, many of them also have other important functions such as pole cell development, neural stem cell maintenance, sex determination, and immune function. Analysis of expression patterns and gene function will be required to determine which of these genes are involved in *D. citri* segmentation, and which might be good targets for control of *D. citri* and Huanglongbing.

## Materials and Methods

*D. citri* genes in genome v3 [24] were identified by BLAST analysis of *D. citri* sequences with insect CHS and UAP orthologs. Reciprocal BLAST of the NCBI non-redundant protein database was used to confirm orthology. Manual annotation of genes was performed in Apollo 2.1.0 using RNA-Seq reads, IsoSeq transcripts and *de* novo-assembled transcripts as evidence. A more detailed description of the annotation workflow is available via protocols.io [25]. Multiple alignments of the predicted *D. citri* proteins and their insect homologs were performed using MUSCLE [26]. Phylogenetic trees were constructed using full-length protein sequences in MEGA7 or MEGAX. Orthologs used in tree construction are listed in Table 3. Gene expression levels were obtained from the Citrusgreening Expression Network [16] and visualized using Excel and the pheatmap package in R [27–28].

**Table 3.**
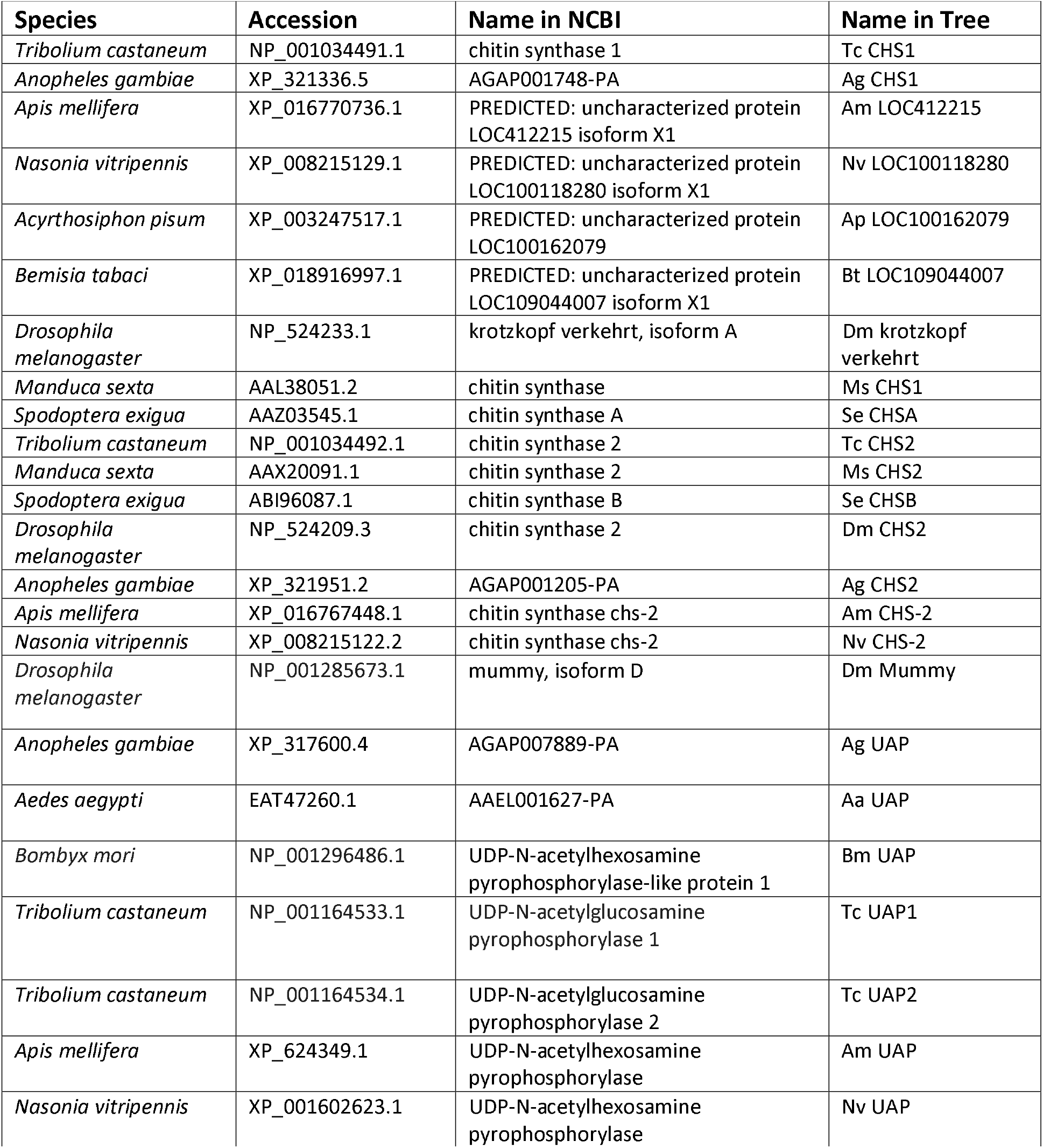
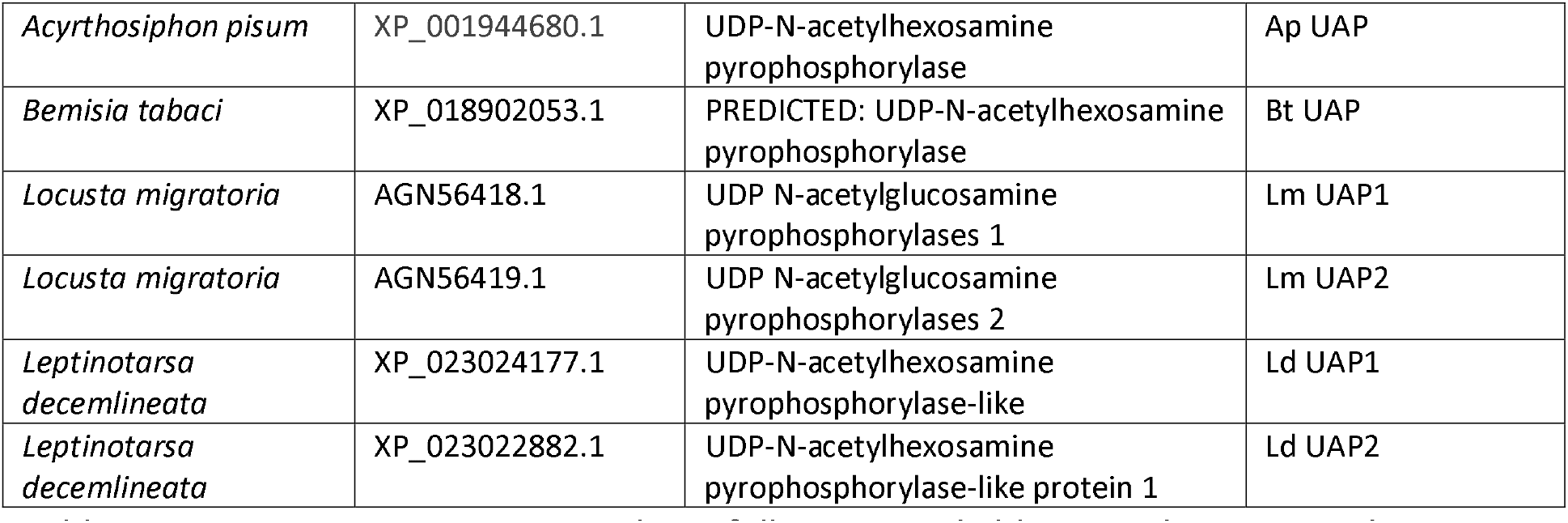
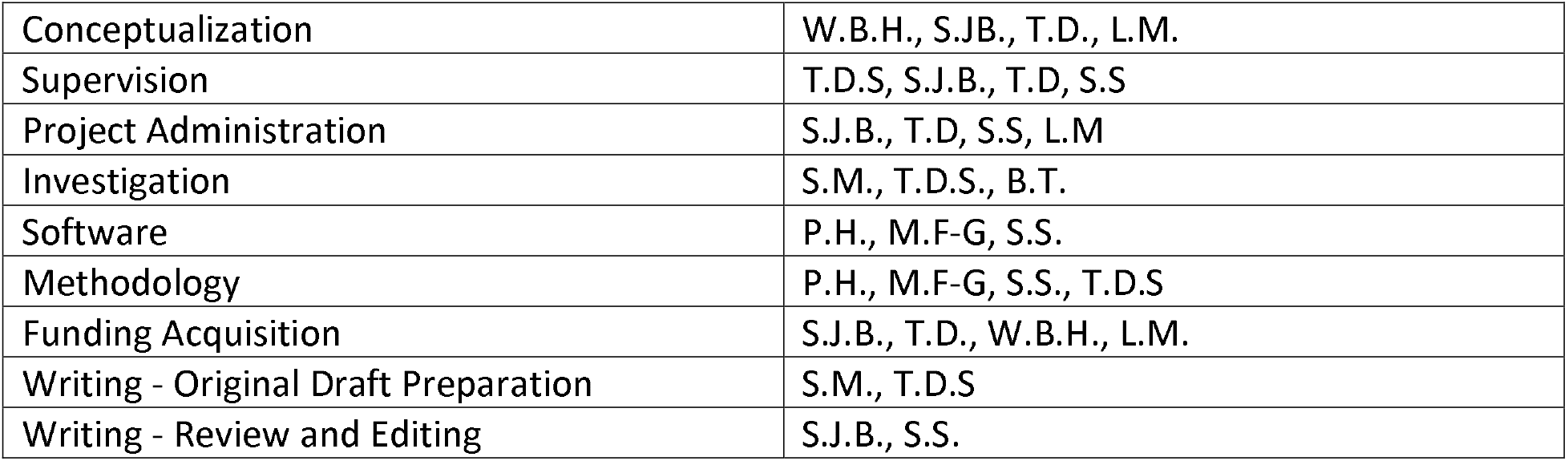
Species, NCBI Accession numbers, full names and abbreviated names used in phylogenetic trees are listed for all orthologs included in phylogenetic analyses (Figure 1,4).

## Acknowledgements

We thank Dr. Josh Benoit for assistance with data visualization. This work was supported by USDA-NIFA grant 2015- 70016-23028 and an Institutional Development Award (IDeA) from the National Institute of General Medical Sciences of the National Institutes of Health under grant number P20GM103418.

## References

1. Zhu KY, Merzendorfer H, Zhang W, Zhang J, Muthukrishnan S. Biosynthesis, turnover, and functions of chitin in insects. Annu Rev Entomol. 2016;61:177–96.

2. Arakane Y, Hogenkamp DG, Zhu YC, Kramer KJ, Specht CA, Beeman RW, et al. Characterization of two chitin synthase genes of the red flour beetle, *Tribolium castaneum*, and alternate exon usage in one of the genes during development. Insect Biochem Mol Biol. 2004;34:291–304.

3. Muthukrishnan S, Merzendorfer H, Arakane Y, Kramer KJ. Chitin metabolism in insects. Insect Mol Biol Biochem. 2012. p. 193–235.

4. Wang Y, Fan H-W, Huang H-J, Xue J, Wu W-J, Bao Y-Y, et al. Chitin synthase 1 gene and its two alternative splicing variants from two sap-sucking insects, *Nilaparvata lugens* and *Laodelphax striatellus* (Hemiptera: Delphacidae). Insect Biochem Mol Biol. 2012;42:637–46.

5. Arakane Y, Muthukrishnan S, Kramer KJ, Specht CA, Tomoyasu Y, Lorenzen MD, et al. The *Tribolium* chitin synthase genes TcCHS1 and TcCHS2 are specialized for synthesis of epidermal cuticle and midgut peritrophic matrix. Insect Mol Biol. 2005;14:453–63.

6. Arakane Y, Specht CA, Kramer KJ, Muthukrishnan S, Beeman RW. Chitin synthases are required for survival, fecundity and egg hatch in the red flour beetle, *Tribolium castaneum*. Insect Biochem Mol Biol. 2008;38:959–62.

7. Zhai Y, Fan X, Yin Z, Yue X, Men X, Zheng L, et al. Identification and functional analysis of chitin synthase A in Oriental Armyworm, *Mythimna separata*. Proteomics. 2017;17.

8. Mohammed AMA, DIab MR, Abdelsattar M, Khalil SMS. Characterization and RNAi-mediated knockdown of chitin synthase A in the potato tuber moth, *Phthorimaea operculella*. Sci Rep; 2017;7.

9. Shi J-F, Mu L-L, Chen X, Guo W-C, Li G-Q. RNA interference of chitin synthase genes inhibits chitin biosynthesis and affects larval performance in *Leptinotarsa decemlineata* (Say). Int J Biol Sci 2016;12:1319–31.

10. Liu X, Cooper AMW, Yu Z, Silver K, Zhang J, Zhu KY. Progress and prospects of arthropod chitin pathways and structures as targets for pest management. Pestic. Biochem. Physiol. 2019. p. 33–46.

11. Silva CP, Silva JR, Vasconcelos FF, Petretski MDA, Damatta RA, Ribeiro AF, et al. Occurrence of midgut perimicrovillar membranes in paraneopteran insect orders with comments on their function and evolutionary significance. Arthropod Struct Dev. 2004;33:139–48.

12. Lu ZJ, Huang YL, Yu HZ, Li NY, Xie YX, Zhang Q, et al. Silencing of the chitin synthase gene is lethal to the asian citrus psyllid, *Diaphorina citri*. Int J Mol Sci. 2019; doi:10.3390/ijms20153734

13. Galdeano DM, Breton MC, Lopes JRS, Falk BW, Machado MA. Oral delivery of doublestranded RNAs induces mortality in nymphs and adults of the Asian citrus psyllid, *Diaphorina citri*. PLoS One 2017;12:e0171847.

14. Zhang X, Zhang J, Park Y, Zhu KY. Identification and characterization of two chitin synthase genes in African malaria mosquito, *Anopheles gambiae*. Insect Biochem Mol Biol. 2012;42:674–82.

15. Hogenkamp DG, Arakane Y, Zimoch L, Merzendorfer H, Kramer KJ, Beeman RW, et al. Chitin synthase genes in *Manduca sexta*: characterization of a gut-specific transcript and differential tissue expression of alternately spliced mRNAs during development. Insect Biochem Mol Biol. 2005;35:529–40.

16. Flores-Gonzalez M, Hosmani P, Fernandez-Pozo N, Mann M, Humann J, Main D, et al. Citrusgreening.org: An open access and integrated systems biology portal for the Huanglongbing (HLB) disease complex. bioRxiv Cold Spring Harbor Laboratory; 2019;868364. doi:10.1101/868364

17. Araújo SJ, Aslam H, Tear G, Casanova J. mummy/cystic encodes an enzyme required for chitin and glycan synthesis, involved in trachea, embryonic cuticle and CNS development - Analysis of its role in *Drosophila* tracheal morphogenesis. Dev Biol. 2005;288:179–93.

18. Tonning A, Helms S, Schwarz H, Uv AE, Moussian B. Hormonal regulation of mummy is needed for apical extracellular matrix formation and epithelial morphogenesis in *Drosophila*. Development; 2006;133:331–41.

19. Schimmelpfeng K, Strunk M, Stork T, Klämbt C. mummy encodes an UDP-N-acetylglucosamine-dipohosphorylase and is required during *Drosophila* dorsal closure and nervous system development. Mech Dev. 2006;123:487–99.

20. Humphreys GB, Jud MC, Monroe KM, Kimball SS, Higley M, Shipley D, et al. Mummy, A UDP-N-acetylglucosamine pyrophosphorylase, modulates DPP signaling in the embryonic epidermis of *Drosophila*. Dev Biol; 2013;381:434–45.

21. Arakane Y, Baguinon MC, Jasrapuria S, Chaudhari S, Doyungan A, Kramer KJ, et al. Both UDP N-acetylglucosamine pyrophosphorylases of *Tribolium castaneum* are critical for molting, survival and fecundity. Insect Biochem Mol Biol. 2011;41:42–50.

22. Liu X, Li F, Li D, Ma E, Zhang W, Zhu KY, et al. Molecular and functional analysis of UDP-N-acetylglucosamine pyrophosphorylases from the migratory locust, *Locusta migratoria*. PLoS One; 2013;8.

23. Shi JF, Fu J, Mu LL, Guo WC, Li GQ. Two *Leptinotarsa* uridine diphosphate N-acetylglucosamine pyrophosphorylases are specialized for chitin synthesis in larval epidermal cuticle and midgut peritrophic matrix. Insect Biochem Mol Biol. 2016;68:1–12.

24. Hosmani P, Flores-Gonzalez M, Shippy T, Vosburg C, Massimino C, Tank W, et al. Chromosomal length reference assembly for *Diaphorina citri* using single-molecule sequencing and Hi-C proximity ligation with manually curated genes in developmental, structural and immune pathways. bioRxiv Cold Spring Harbor Laboratory; 2019;869685. doi:10.1101/869685

25. Shippy, TD; Miller, S; Massimino, C; Vosburg, C; Hosmani, PS; Flores-Gonzalez, M; Mueller, LA; Hunter, WB; Benoit, JB; Brown, SJ; D’elia, T; Saha S. Annotating genes in Diaphorina citri genome version 3. protocols.io. 2020; doi:10.17504/protocols.io.bniimcce

26. MUSCLE. https://www.ebi.ac.uk/Tools/msa/muscle/. Accessed Oct 26 2020.

27. R Core Team. R: A language and environment for statistical computing. R Foundation for Statistical Computing, Vienna, Austria. 2020. URL https://www.R-project.org/. Accessed Dec 17 2020.

28. Kolde R. pheatmap: Pretty Heatmaps. R package version 1.0.12. 2020.

29. Adams MD, Celniker SE, Holt RA, Evans CA, Gocayne JD, Amanatides PG, et al. The genome sequence of *Drosophila melanogaster*. Science. 2000;2185–95.

30. Holt RA, Mani Subramanian G, Halpern A, Sutton GG, Charlab R, Nusskern DR, et al. The genome sequence of the malaria mosquito *Anopheles gambiae*. Science. 2002;298:129–49.

31. Matthews BJ, Dudchenko O, Kingan SB, Koren S, Antoshechkin I, Crawford JE, et al. Improved reference genome of *Aedes aegypti* informs arbovirus vector control. Nature. 2018;563:501–7.

32. Tribolium Genome Sequencing Consortium, Richards S, Gibbs RA, Weinstock GM, Brown SJ, Denell R, et al. The genome of the model beetle and pest *Tribolium castaneum*. Nature. 2008;452:949–55.

33. Elsik CG, Worley KC, Bennett AK, Beye M, Camara F, Childers CP, et al. Finding the missing honey bee genes: lessons learned from a genome upgrade. BMC Genomics. 2014; doi:10.1186/1471-2164-15-86.

34. Werren JH, Richards S, Desjardins CA, Niehuis O, Gadau J, Colbourne JK, et al. Functional and evolutionary insights from the genomes of three parasitoid *Nasonia* species. Science. 2010; doi:10.1126/science.1178028.

35. International Aphid Genomics Consortium. Genome sequence of the pea aphid *Acyrthosiphon pisum*. PLoS Biol. 2010; doi:10.1371/journal.pbio.1000313

36. Chen W, Hasegawa DK, Kaur N, Kliot A, Pinheiro PV, Luan J, et al. The draft genome of whitefly *Bemisia tabaci* MEAM1, a global crop pest, provides novel insights into virus transmission, host adaptation, and insecticide resistance. BMC Biol. 2016. doi:10.1186/s12915-016-0321-y.

37. Wu Z, Zhang H, Bin S, Chen L, Han Q, Lin J. Antennal and abdominal transcriptomes reveal chemosensory genes in the Asian Citrus Psyllid, *Diaphorina citri*. PLoS One; 2016;11.

